# Financial incentives facilitate the neural computation of prosocial decisions stronger in low empathic individuals

**DOI:** 10.1101/2021.03.23.436445

**Authors:** Vassil Iotzov, Anne Saulin, Jochen Kaiser, Shihui Han, Grit Hein

**Affiliations:** Translational Social Neuroscience Lab, Department of Psychiatry, Psychosomatic and Psychotherapy, University Hospital of Wuerzburg, Margarete-Höppel-Platz 1, 97080 Würzburg / Germany; Institute of Medical Psychology, Goethe-Universität, Heinrich-Hoffmann-Str. 10, Building 93A, Room A45, 60528 Frankfurt am Main; Culture and Social Cognitive Neuroscience Lab, School of Psychological and Cognitive Sciences, PKU-IDG/McGovern Institute for Brain Research, Peking University, Room 1614, Wang Kezhen Building, No. 52, Haidian Road, Haidian District, Beijing 100080, People’s Republic of China

**Keywords:** Empathy, prosocial behaviour, incentives, drift-diffusion modelling, fMRI

## Abstract

Financial incentives are commonly used to motivate behaviours. There is also evidence that incentives can decline the behaviour they are supposed to foster, for example, documented by a decrease in blood donations if a financial incentive is offered. Based on these findings, previous studies assumed that prosocial motivation is shaped by incentives. However, so far, there is no direct evidence showing an interaction between financial incentives and a specific prosocial motive. Combining drift-diffusion modelling and fMRI, we investigated the effect of financial incentives on empathy, i.e., one of the key motives driving prosocial decisions. In the empathy-alone condition, participants made prosocial decisions based on empathy, in the empathy-bonus condition, they were offered a financial bonus for prosocial decisions, in addition to empathy induction. On average, the bonus enhanced the information accumulation in empathy-based decision. On the neural level, this enhancement was related to the anterior insula, the same region that also correlated with empathy ratings. Moreover, the effect of the financial incentive on anterior insula activation was stronger the lower a person scored on empathy. These findings show that financial incentives enhance prosocial motivation in the absence of empathy but have little effect on high empathic individuals.

## Introduction

Financial incentives are frequently used to motivate people. Such measures are based on empirical evidence showing that financial incentives increase the frequency of the rewarded behaviour (Garbers and Konradt, 2014; Wei and Yazdanifard, 2014), including cooperative and prosocial behaviours (Balliet *et al*., 2011; Stoop *et al*., 2018). For example, in a meta-analysis, Balliet and colleagues found that reward positively affects cooperation (Balliet *et al*., 2011). Consequently, financial incentives could increase the motivation to behave prosocially (Ariely *et al*., 2009). However, there is other evidence that incentives can undermine the very behaviour they are meant to strengthen (Titmuss, 1970; Deci *et al*., 1999; Benabou and Tirole, 2006; Murayama *et al*., 2010; Niza *et al*., 2013; Rode *et al*., 2015; Besley and Ghatak, 2018). The most classic example in the realm of prosocial behaviours is the observation that people donate less blood if they are paid to do so, compared to the amount of blood that they donate without payment, i.e., to help others (Titmuss, 1970; Niza *et al*., 2013). In line with these observations, other studies have shown that adding financial incentives can reduce prosocial behaviours (Bowles, 2008; Ariely *et al*., 2009; Holmås *et al*., 2010). In sum, the evidence regarding the effects of incentives on prosocial decisions is inconsistent, and mainly based on behavioral observations that do not provide insights in the underlying motivational processes. As a result, it remained unclear whether and how financial incentives interact with a specific prosocial motive.

Overcoming this limitation, our study directly investigated how a financial incentive shapes prosocial decisions that are driven by a specific prosocial motive, i.e., empathy. Incorporating previous approaches, we used a well-established decisions task (i.e., a modified version of a binary dictator game (Hein *et al*., 2016b)). Extending previous studies, we activated a specific prosocial motive (empathy) before participants entered the decision task, and, in one condition, added a financial incentive. This allowed us to investigate how financial incentives change the processing of prosocial decisions that are driven by one specific, carefully controlled motive. To control for other motivations that might play a role besides empathy (self-image concerns; reciprocity), the incentive was offered in private, the decisions were kept anonymous, and the participants knew that they would not meet the other players after the study. This measure is important because it minimizes participants’ motivation to maintain a positive public image, i.e., a different motive that may affect participants’ prosocial decisions besides empathy (Benabou and Tirole, 2006; Ariely *et al*., 2009; Exley, 2017; Besley and Ghatak, 2018).

Empathy is defined as the affective response to another person’s misfortune (Batson *et al*., 1995; Lamm *et al*., 2011; Decety *et al*., 2016; Hein *et al*., 2016b; Marsh, 2018). Neuroscientific studies have shown that prosocial decisions correlate with brain activations in regions that are also associated with individual differences in empathy, such as the anterior insula (AI) cortex and the anterior cingulate cortex (ACC) (Hein *et al*., 2010; Masten *et al*., 2011; Hein *et al*., 2016b; Marsh, 2018). We chose to induce empathy because it is one of the strongest prosocial motives (Batson *et al*., 1995; Decety *et al*., 2016). Previous work has established a reliable link between the individual strength of the empathy motive and the propensity to act prosocially, e.g., decisions that maximize the outcome of another person at costs to oneself (Batson *et al*., 1995; Decety *et al*., 2016). The stronger the empathy motive, the stronger the propensity to decide in favour of the other person. Previous social psychology work has investigated how empathy is shaped by selfish motives, such as the motive to withdraw from a stress-inducing situation (Batson *et al*., 1981). However, to the best of our knowledge, there are no previous studies that tested how financial incentives affect the components of empathy-based prosocial decisions.

The study consisted of two parts (**Fig. 1**). In part 1, the empathy motive was activated towards one partner (a confederate). In the following allocation task, participants allocated points to the respective partner (here driven by empathy; empathy-alone condition). Next, the confederate was replaced by a new individual that served as a partner for part 2. In part 2, the empathy motive was activated again. However, before starting the decision task, the participant was told that she would receive a bonus if she decided prosocially in the majority of the decision trials. In the following allocation task, participants again allocated points to the respective partner (here driven by empathy and the financial incentive; empathy-bonus condition). The order of the two conditions (empathy-alone and empathy-bonus) was counterbalanced across participants and the two confederates.

**Fig. 1.**
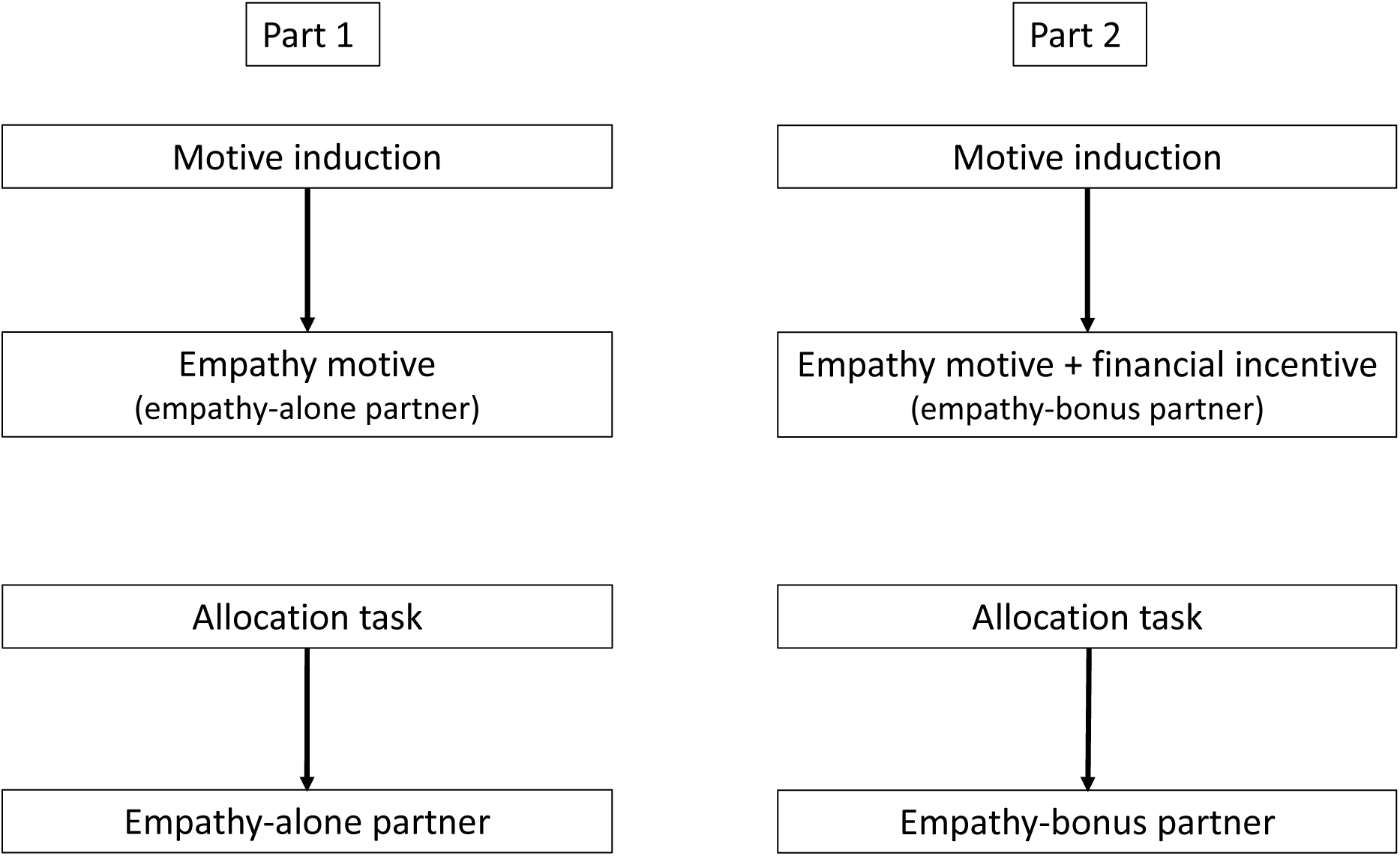
Overview of an exemplary experimental procedure. The study consisted of two parts. In this example, in part 1, the empathy motive was activated towards one confederate (the empathy-alone partner). In the following allocation task, participants allocated points to the empathy partner (i.e., driven by the empathy motive). Next, the confederate was replaced by a new individual that served as partner for part 2. Again, the empathy motive was activated. After the empathy motive induction additionally a bonus for choosing the prosocial option in the majority of trials in the subsequent allocation task was offered (empathy-bonus partner). Thus, in the following allocation task, participants allocated points towards the empathy-bonus partner (i.e., driven by the empathy motive and the additionally offered bonus). The order of motive induction (empathy-alone, empathy-bonus) was counterbalanced across participants and both confederates. The respective partner was indicated by a cue in one of two counterbalanced colors.

To induce empathy, participants repeatedly observed two interaction partners receiving painful shocks in a number of trials, a situation known to elicit an empathic response (Lamm *et al*., 2011; Hein *et al*., 2016a; Hein *et al*., 2016b). As a measure of the individual strength of the induced empathy motive, participants rated how they felt when observing the respective other person in pain (**Fig. 2A**). To allow participants to simulate the state (pain) of the other person, in some trials, participants received painful stimulation themselves.

**Fig. 2.**
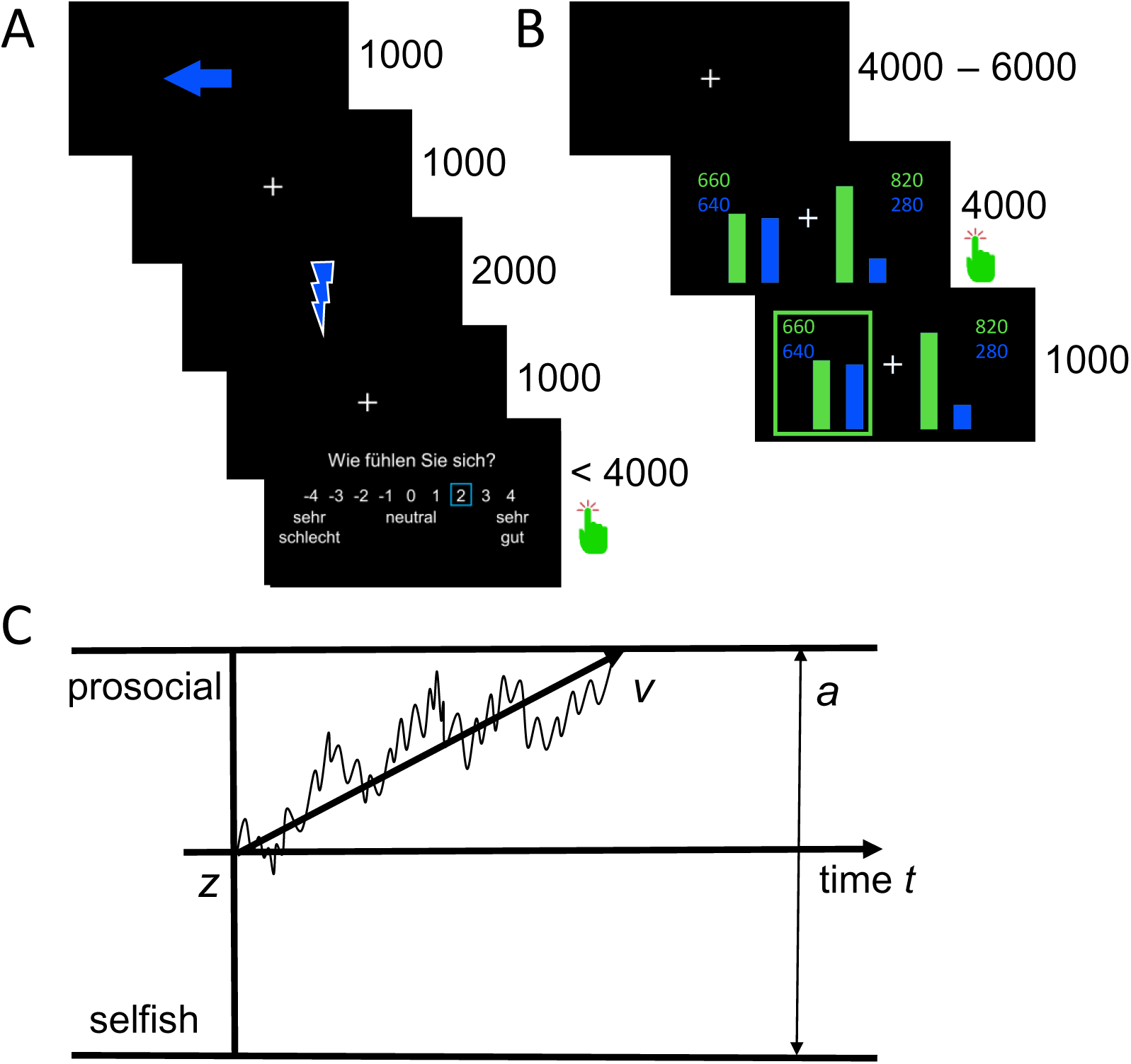
Examples of induction and decision trials and schematic overview of the drift-diffusion model (DDM). **A)** Example trial of the empathy induction. The arrow cue indicated the receiver of the stimulation (self, the empathy-alone partner in one condition or the empathy-bonus partner in the other condition). The lightning bolt indicated pain stimulation. Participants rated how they felt after observing the stimulation of the partner or receiving it themselves (-4 = very bad; +4 = very good). **B)** Example trial of the allocation task. Participants chose between a prosocial option that maximized points for the partner or a selfish option that maximized points for themselves. In this example trial, the participant chose the prosocial option, which maximized the outcome of the partner at a cost to the participant (green box). **C)** Schematic overview of the drift-diffusion model. According to the drift-diffusion model, the decision process is a noisy accumulation of information (jagged black line). From the distributions of both prosocial and selfish decisions, a set of parameters is estimated that allows to draw conclusions about the underlying cognitive processes. These are mainly the speed of information accumulation (*v*-parameter), the starting point of the decision process (*z*-parameter), and the amount of information to be processed (*a*-parameter). As soon as the accumulated information reaches one of the two boundaries, the decision is made (upper boundary = prosocial option; lower boundary = selfish option).

During the allocation task inside the fMRI scanner, participants allocated points to the partners at a cost to themselves (**Fig. 2B**). The allocation of points towards the one partner (empathy partner) should be based on the previously activated empathy motive (empathy-alone condition). The allocation of points towards the other partner (empathy-bonus partner) was also based on the previously activated empathy motive. However, in this condition, participants were additionally informed that they would receive a bonus for choosing the prosocial option in the majority of trials in the subsequent allocation task (empathy-bonus condition). Note that the bonus corresponded to the maximally possible outcome in the allocation task (i.e., the outcome that a participant would gain if she always chose the selfish option). Thus, deciding prosocially to reach the bonus criterion in the empathy-bonus condition did not result in a financial loss for the participants.

To specify how incentives modulate empathy-related decisions, we used drift-diffusion modelling (DDM). DDMs assumes that during binary decisions, noisy information is accumulated to select a decision option mainly based on three different parameters (the *v-*, *z-* and *a*-parameters; (Forstmann *et al*., 2016; Ratcliff *et al*., 2016); **Fig. 2C**). The *v*-parameter describes the speed of noisy evidence accumulation in order to choose one of two options, i.e., the efficiency of the decision process itself. Thus, a larger *v*-parameter indicates faster information accumulation regarding the prosocial option. The individual decision bias is reflected by the *z*-parameter. In contrast to the *v*-parameter, the *z*-parameter models the individual preferences with which a person starts the decision process. For example, if a person has a strong prior preference for prosocial decisions, the starting point of the decision process is closer to the prosocial decision boundary, and therefore less evidence has to be accumulated regarding the prosocial option. The amount of evidence that needs to be accumulated to distinguish between the two options is reflected by the *a*-parameter. We modelled these three parameters (*v, z*, and *a*) for decisions that were driven by the empathy motive alone and that were driven by the combination of the empathy motive and the financial incentive, based on the raw data from the entire data set (i.e., including trial-by-trial information of all decisions). Additionally, the non-decision time (*t_0_*) was estimated across conditions (see Methods for details).

Extending the classical DDM approach, a recent model has proposed that the evidence in favour of one or another choice alternative might be shaped by affective and motivational states (Roberts and Hutcherson, 2019). Supporting this assumption, affective states have been found to change central parameter of the choice process such as the drift rate (*v*-parameter; (Lerche *et al*., 2018; Roberts and Hutcherson, 2019; Aylward *et al*., 2020; Thompson and Steinbeis, 2021)) and the starting point (*z*-parameter, (White *et al*., 2018)). Inspired by these results, we assumed that the evidence in favor of a prosocial choice might be different in different motivational states (i.e., induced by empathy and its potential interaction with the incentive), reflected by a changes in the drift rate and/ or the starting point.

One assumption is that financial incentives may enhance empathy-related prosocial decisions, inspired by findings of reward-related increases of prosociality (Garbers and Konradt, 2014; Wei and Yazdanifard, 2014). If this was true, the frequency and efficiency of prosocial decisions should be higher in the empathy-bonus compared to the empathy-alone condition. Specifying the potential effect of the incentive on the prosocial choice process, the DDM proposes that an incentive-related facilitation of prosocial choices may originate A) from an increased speed of information accumulation, i.e., an increased drift rate (*v*-parameter (Lerche *et al*., 2018; Roberts and Hutcherson, 2019; Aylward *et al*., 2020; Thompson and Steinbeis, 2021), an enhancement of participants’ initial preference to choose the prosocial option, i.e, a shift of the starting point towards the prosocial decision boundary (*z*-parameter (White *et al*., 2018)), or from an enhancement of the *v*- as well as the *z*-parameter in the empathy-bonus compared to the empathy-alone condition,

Alternatively, it is possible that financial incentives undermine empathy-related prosocial decisions, in line with previous findings that showed an incentive-related decrease in prosocial behaviour (Titmuss, 1970; Benabou and Tirole, 2006; Murayama *et al*., 2010; Rode *et al*., 2015). In this case, prosocial decisions should be more frequent in the empathy-alone compared to the empathy-bonus condition. According to the DDM, such an undermining effect may be reflected A) by a reduced speed of information accumulation (*v*-parameter), B) a shift of the starting point away from the prosocial decision boundary (*z*-parameter), or C) a reduction in both parameters in the empathy-bonus compared to the empathy-alone condition.

Finally, it is possible that the effect of financial incentives depends on the strength of the empathy motive, i.e., might be different for high empathic compared to low empathic individuals. If this is true, the individual difference between the empathy-bonus vs empathy-alone condition and changes in the drift rate and/or the starting point should be related to the individual empathy ratings, i.e., the measure that captures the strength of the empathy motive during the first part of the study.

Based on previous evidence that has linked empathy-related decisions to neural responses in the AI (Hein *et al*., 2010; Masten *et al*., 2011; Hein *et al*., 2016b; Marsh, 2018), we assume that an incentive-related increase in the *v*- and/or *z*-parameter (reflecting facilitation of empathy-related decisions) is associated with an increase in brain regions associated with the processing of empathy and empathy-related decisions such as the AI and the ACC. In contrast, an incentive-related decrease in the *v*- and/or *z*-parameter (reflecting a potential undermining effect) should be related to a decrease in AI and ACC activity.

## Methods

### Materials and Methods

#### Participant details

33 healthy women (mean age was 25.05 years, *s.e.* = 0.74) participated in the study. We chose a female instead of a gender-mixed subject group because it allowed us to choose female confederates and thus to avoid the potential complications of gender-mixed pairing of participants and confederates. The confederates were two female students, trained to play their roles in counterbalanced order. The data from two participants had to be discarded as outlier (frequency of prosocial decisions, 3.42 SDs below the mean (*M_empathy-alone_* = 44.35, *SD_empathy-alone_* = 12.97). Thus, we analyzed 31 data sets. We obtained ethics approval (EK 458122014) for conducting the study and written informed consent from our participants. The experiment was conducted following the Helsinki guidelines. Participants received monetary compensation (show up fee plus payout from two randomly chosen trials of the allocation task; see below).

### Procedure

#### Overall procedure

Outside the fMRI scanner, we attached pain electrodes to the back of the participants’ and the confederates’ hands and determined the individual thresholds for painful and painless stimulation using a standard procedure (Hein *et al*., 2016a; Hein *et al*., 2016b). Next, the participant and the confederates played a manipulated lottery (drawing matches) that ostensibly determined the amount of pain the person would receive in the following task. Because the empathy induction required saliently more pain for the confederates, the drawing of the matches was organized in such a way that the participant always drew the last match and thus was assigned to receive only a few painful stimuli.

The participant was placed inside the fMRI scanner, and one of the confederates was placed on a chair next to the participant in the scanner room. The confederate’s hand with the pain electrode was placed on a tilted table over the participants’ knee. Through a mirror in the head coil, participants could see the hand of the other, together with the visual stimulation on a screen that was positioned at the end of the fMRI bed. During the empathy induction, participants either saw a dark-coloured flash (painful stimulation) or a light-coloured flash (non-painful stimulation), indicating the intensity of the stimulation of the confederate. In a small portion of trials (five from fifteen), they received pain stimulation themselves, indicated by a dark-coloured flash of a different colour. During the decision task, participants were presented two options to allocate points between themselves and the other person. Colours were counterbalanced across participants.

The study started with the empathy induction, followed by the allocation task towards the first confederate. After replacing this confederate, the same procedure (empathy induction followed by the allocation task) was repeated with the second confederate (**Fig. 1**). In the empathy-alone condition, the allocation task started immediately after the empathy induction. In the empathy-bonus condition, after the empathy induction, participants were told that they would receive a bonus (additional 5 Euro) if they chose the prosocial option in the majority of trials. We deliberately refrained from specifying the percentage of prosocial decisions that were required to win the bonus to avoid strategy effects. However, participants knew that the bonus would compensate the maximally possible outcome in the allocation task (i.e., the outcome that a participant would gain if she always chose the selfish option). Thus, deciding prosocially to reach the bonus in the empathy-bonus condition did not result in a financial loss for the participants. To minimize reputation effects, participants received the bonus information in private without the partner’s knowledge.

Apart from the bonus in the empathy-bonus condition, the experimental procedure was identical in both conditions. The order of the conditions and the assignment of the confederates was counterbalanced across participants. At the end of the experiment, both confederates left, and the participants stayed in the scanner until anatomical image acquisition was completed. Finally, participants were asked to complete the Interpersonal Reactivity Index (IRI; (Davis, 1980)), and a scale that assessed their impression of both confederates (Hein *et al*., 2016a). The impression ratings were comparable between confederates (lmm *χ^2^_(1)_* = 0.36, *p* = .55, *B* = -0.10, *s.e.* = 0.16).

Participants spent approximately 60 min inside the scanner, and the entire procedure lasted about 2 hours. In addition to the show-up fee, participants received the payout from two randomly chosen allocation trials, and the bonus of five Euro if they had made prosocial decisions in 75% of all trials.

All ratings during the induction phase and all decisions in the allocation task were kept anonymous. Particular care was taken to ensure that this was clear to participants by pointing out the following: Inside the scanner room, the partner had a separate visual display, such that the participant viewed stimuli via back-projection from a mirror onto a screen, while the confederates beside the scanner viewed stimuli via cardboards/video glasses with a built-in display (Hein *et al*., 2016a). Thus, all ratings and decisions were private and could not be observed by the other participants (Hein *et al*., 2016a). Moreover, participants knew that they would not meet after the experiment because the scanned participant needed to stay longer for an anatomical scan. The experimenter was outside the scanner room, and it was pointed out that he could not see the ratings and decisions either.

#### Empathy induction

In each empathy-induction trial, first we presented a coloured arrow indicating the person who will receive the following electric stimulation for 1000 ms. After this cue, a fixation cross was presented for 1000 ms, followed by a coloured lightning bolt shown for 2000 ms. Participants were informed that a blinking dark-coloured lightning bolt indicates a painful stimulus, whereas a blinking light-coloured lightning bolt indicates a non-painful stimulus. After receiving or observing the electric stimulation, we showed a 9-point rating scale with the question “How do you feel?”. The scale ranged from -4 (labeled “*very bad*”) to +4 (labeled “*very good*”). Participants had to respond within 4000 ms (**Fig. 2A**). The empathy induction consisted of 30 trials: 10 that were ostensibly painful for the partner (other-pain trials), 5 that were not painful for the partner (other-no-pain trials), 5 painful trials for the participant (self-pain trials), and 10 non-painful trials (self-no-pain trials) for the participant. The self-pain trials were added to allow participants to simulate the state (pain) of the other person. To test their potential influence on empathy changes, we compared the ratings in other-pain trials that were preceded by a self-pain trial (i.e., empathy ratings under the condition of self-pain experience) with the ratings in other-pain trials that were preceded by an other-pain trial (i.e., empathy ratings without preceding self-pain experience). The results showed no difference between the other-pain ratings after self-pain and the other-pain ratings without prior self-pain (t_(61)_ = 0.34, p = .73). Based on these results, the self-pain experience had no significant effect on empathy changes during empathy induction.

#### Allocation task

The allocation task was identical in both conditions and based on a well-established paradigm (Hein *et al*., 2016b). In each trial, participants allocated points to themselves and the respective partner (**Fig. 2B**) and could choose between maximizing the relative outcome of the other person by reducing their own relative outcome (prosocial choice) and maximizing their own relative outcome at a cost to the partner (selfish choice). The outcome was relative to the outcome that the participant would have gained when choosing the other option. The initial number of points was always higher for the participant compared to the partners. This measure was inspired by previous behavioral economics research, showing that participants make more prosocial decisions if their initial payoff is higher than the partner’s payoff (“advantageous inequality” (Fehr and Schmidt, 1999; Bolton and Ockenfels, 2000; Charness and Rabin, 2002)). The choice options used in the present study created advantageous inequality to optimize the number of prosocial choices, which was the main focus of our study.

For the point distributions, we used values between 900 and 1200. The respective value was divided into a self:other ratio of 60:40 or of 90:10. Each trial of the allocation task contained a prosocial and a selfish option. The prosocial option was always the more egalitarian option, with a point distribution of 60% (self) to 40% (other). In contrast, in the selfish option, points were allocated with a ratio of 90% (self) to 10% (other). Participants’ losses were symmetrical to the partner’s gains. For example, a total of 1000 points were distributed with self:other ratios of 60:40 (600:400 points), 90:10 (900:100 points). Thus, the participant’s loss is 900 - 600 = 300 points, which corresponds exactly to the gain of the partner (400 - 100 = 300 points). We used these fixed and symmetrical ratios to minimize unspecific effects of loss aversion.

Each decision-trial started with an inter-trial interval indicated by a fixation cross presented for a period jittered between 4000 and 6000 ms (**Fig. 2B**). After this, participants saw the two possible distributions of points in different colours, indicating the potential gain for the participant and the potential gain for the current partner. Participants had to choose one of two distributions within 4000 ms by pressing the left button on a response box to select the distribution on the left side and the right button to select the distribution on the right side. The position of the two allocation options was randomized across trials to minimize response biases due to motor habituation. A green box appeared around the distribution that was selected by the participant at 4000 ms after distribution onset. The box was shown for 1000 ms. At the end of the experiment, two of the distributions chosen by the participant were randomly selected for payment (100 points = 50 cents). Participants performed 60 decision trials in each motive-induction condition, i.e., 120 trials in total.

#### Pain stimulator

For pain stimulation, we used electrical stimulation (bipolar, monophasic; output range 5Hz, 0-10 mA) from a single-current stimulator (Neurometer CPT/C; Neurotron Inc.). After attaching the electrodes at the index finger of the right hand and connecting them to the single-current stimulator, the respective person was asked to press the button for defining the current threshold and to decide when she is feeling the stimulation – the value of this threshold was used as painless stimulation. In a second run the participant was asked to press the same button, but now to hold it pressed until the pain was at an unacceptable level and then to release – this threshold was used for the painful stimulation.

### Experimental design and statistical analyses

The aim of our study was to compare prosocial decisions driven by empathy alone with prosocial decisions driven by a combination of empathy and a financial bonus. Therefore, we used a within-subject design in which each participant performed the identical social decision task under two different conditions: the empathy-bonus and the empathy-alone condition. Behavioural data were analyzed with R-Studio Version 1.1.463(RStudio Team, 2020) and R Version 3.6.0(RCore Team, 2019) and Python (HDDM; Spyder Version 3.3.2; Python Version 2.7.15 (Van Rossum, 2007; Wiecki *et al*., 2013)).

#### Regression analyses

All regression analyses were performed with the R-packages “stats” (RCore Team, 2019) using, “lme4” (Bates *et al*., 2015), “car” (Fox and Weisberg, 2019), and MuMIn (Bartoń, 2019). Results were visualized with the “tidyverse” package (Wickham *et al*., 2019). All continuous predictors in our regressions are z-scored.

Empathy ratings showed a right-skewed distribution (Shapiro-Wilk *W* = .94, *p* < .01), so the data was log-transformed to normal distribution. Pearson correlation was computed between the empathy ratings and the empathic concern scale (EC) from the Interpersonal Reactivity Index (IRI)(Davis, 1980). In further data analyses, we used linear models within condition and linear mixed models (lmm) with participants as random effect between conditions.

#### Drift-Diffusion Modeling

We choose the DDM, because of its small but trackable number of key parameters and because it is relatively easy to reduce other sequential sampling models (SSMs) to the DDM given specific parameter constraints (Bogacz *et al*., 2006). Moreover, because of the increasing popularity of DDMs in psychology research, the DDM results from our study can be embedded in the existing literature. We used hierarchical drift-diffusion modelling (HDDM (Vandekerckhove *et al*., 2011; Wiecki *et al*., 2013)), which is a version of the classical drift-diffusion model that exploits between-subject and within-subject variability using Bayesian parameter estimation methods and thus is ideal for use with relatively small sample sizes. The analyses were conducted using the python implementation of HDDM (Wiecki *et al*., 2013). Based on previous studies showing changes in drift rate (Lerche *et al*., 2018; Roberts and Hutcherson, 2019; Aylward *et al*., 2020; Thompson and Steinbeis, 2021) and the starting point (White *et al*., 2018) if decisions are made in different affective states, we assumed that these two parameters might also be affected by motivational states. However, given that the modulation of affect and motivation is not the same, effects on the third parameter (the *a*-parameter) are also possible. Therefore, we estimated the full model with *v*, *z*, and *a* possibly being modulated by our two conditions. Moreover, we estimated the non-decision parameter (*t_0_*), which indicates the duration of all extradecisional processes like basic encoding or motor processes (Voss *et al*., 2004). In paradigms like ours that used an identical experimental setting across conditions, it was recommended to estimate the *t_0_*-parameter across conditions (Wagenmakers *et al*., 2008; Servant *et al*., 2014; Nunez *et al*., 2017). Following this recommendation, we estimated the *t_0_*-parameter across the empathy-bonus and the empathy-alone conditions (mean *t_0_* = 0.58, s.e. = 0.02), and refrained from estimating it for each condition separately (see full HDDM results table at github.com (https://github.com/Vassil-Iotzov/empathy_incentives)).

We conducted the same DDM analyses with two different inputs. In one analysis, the input of the DDM was defined categorically based on the type of response (1 = prosocial option; 0 = selfish option). In the other analyses, we used the trial-by-trial point difference (self-loss or other-gain) as additional covariate effecting the drift rate to estimate a hierarchical random intercept model (see Chen and Krajbich (2018) for a similar approach). Other input parameters were reaction time (in seconds), condition (empathy-bonus, empathy-alone), and participants number (0 to 30).

To evaluate the model fit, we conducted posterior predictive checks by comparing the observed data with 500 datasets simulated by our model, thus using the method that has been particularly recommended for HDDMs to obtain quantile comparison and 95% credibility (Wiecki *et al*., 2013)). The respective quantile comparison table is provided at github.com (https://github.com/Vassil-Iotzov/empathy_incentives). Moreover, model convergence was checked by visual inspection of the estimation chain of the posteriors, as well as computing the Gelman-Rubin Geweke statistic for convergence (all values < 1.01) (Gelman and Rubin, 1992). Parameters of interest from the model were extracted for further analysis. Specifically, for each participant, the condition-specific *v*-parameters, *z*-parameters, and *a*-parameters were extracted (resulting in 6 parameters per participant). For the parameter comparison, the posteriors were analyzed directly, as recommended by Wiecki *et al*. (2013). Specifically, the probability was tested that the *v-*, *z-* or *a-*parameter was greater in the empathy-bonus compared to the empathy-alone condition.

The DDM results were visualized using a custom-made R-function based on ggplot2 (part of the “tidyverse”-Package; (Wickham *et al*., 2019)). The following equation was used to calculate the slopes of the *v*-parameters (Alexandrowicz, 2018):

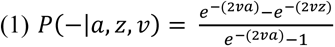

The equation was simplified by setting the variance of the Brownian motion at *s²* = 1 (Alexandrowicz, 2018) in the basic formula:

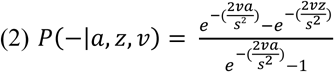

The *a*-parameter was displayed by taking the higher alpha as 100% and calculating the lower alpha according to the respective ratio. The *z*-parameter was plotted as relative *z* (*zr*) also in relation to the *a*-parameter. The full script is available at github.com (https://github.com/Vassil-Iotzov/ggddm).

#### Image Acquisition and Analyses

The experiment was conducted on a 3-T Siemens Magnetom Prisma whole-body MR scanner (Siemens Healthineers), equipped with a one-channel Siemens head coil. Scanner noise was reduced with soft foam earplugs, and head motion was minimized with foam pads. Stimuli presented in the induction phase and in the allocation task were projected onto a rear projection screen located in the front of the scanner. Behavioural responses were recorded with a five-key fibre-optic response box placed on the right hand, and when necessary, vision was corrected using MRI-compatible lenses that matched the dioptre of the participant. Structural image acquisition consisted of 176 T1-weighted transversal images (voxel size of 1 mm) (Hein *et al*., 2016a). Functional imaging data was collected during the allocation task, using T2*-weighted echo-planar imaging (32 slices, slice thickness of 3 mm, ascending acquisition; repetition time, 2100 ms; echo time, 30 ms; flip angle, 80°; field of view, 240 mm; matrix, 80 × 80). In every decision session, 300 images were acquired - a total of 600 Images for both sessions.

#### Preprocessing and statistical model

The images were analyzed with SPM12 (Functional Imaging Laboratory, 2019) and Matlab version 8.6 (Matlab, 2015). Images were preprocessed following the standard procedure recommended in the SPM manual (Functional Imaging Laboratory, 2019), including realignment, slice time correction, coregistration, segmentation, normalize, smoothing.

First-level analyses were performed with the general linear model (GLM), using a canonical hemodynamic response function (HRF). For each of the conditions (empathy-alone and empathy-bonus condition), the respective regressors of prosocial choice trials were included as regressors of interest. The prosocial decisions regressor spanned the period from the onset of the decision screen until the participants’ reaction (average of 1146.37 ms). Regressors of no interest included the period from the participants’ reaction to decision offset (average of 2853.63 ms) and the immediately following period showing the participants’ decision (1000 ms).

Sixteen of our participants made less than five selfish decisions in at least one condition. To avoid empty cells in the model, we refrained from computing direct contrasts between prosocial and selfish choices, and selfish choices were included as regressor of no interest.

For the second-level analyses, contrast images for comparisons of interest (empathy-bonus > implicit baseline, empathy-alone > implicit baseline, empathy-bonus > empathy-alone, and empathy-alone > empathy-bonus) were initially computed on a single-subject level. In the next step, the individual images of the main contrast of interest (empathy-bonus > implicit baseline) were regressed against the *v*-parameter. Results were thresholded using 5% family wise error (FWE) corrected voxel-based inference. We also conducted exploratory analyses using 5% FWE cluster-based inference with a cluster-forming threshold of P_uncorrected_ < .001 and a minimal cluster size of k = 50 and used this threshold for the visualization of our results. Beta estimates were extracted from the entire clusters of activation in the anterior insula obtained from 5% FWE cluster-based inference with P < .001 cluster-forming threshold, k = 50, using MarsBaR (Brett, 2002). Moreover, the respective beta-estimates were extracted from an independent region of interest, defined based on a 20mm sphere around the peak coordinates (x = -43; y = 14; z = 7) from a significant activation likelihood cluster found across all pain empathy experiments in a current meta-analysis (Jauniaux *et al*., 2019).

#### Code and data availability

Behavioural data and scripts are available at github.com (https://github.com/Vassil-Iotzov/empathy_incentives). Imaging data are available at neurovault.org (https://identifiers.org/neurovault.collection:7568).

## Results

### Empathy was induced with comparable strength in both conditions

To quantify the strength of the induced empathy, we calculated the participants’ trial-by-trial ratings while observing the partner in pain relative to their self-pain ratings. Comparing the ratings between the empathy-alone and the empathy-bonus condition revealed no significant differences between conditions (lmm *χ^2^_(1)_* = 0.0001, *p* < .99, *B* = -0.002, *s.e.* = 0.22, *R²_m_* < .01), indicating that empathy was induced with comparable strength in the empathy-alone and the empathy-bonus condition.

The average of the individual empathy ratings in both conditions, i.e., our measure of state empathy, correlated significantly with individual differences in trait empathy assessed with the empathic concern scale (EC) of the Interpersonal Reactivity Index (IRI; (Davis, 1980)), *r*_(29)_ =.36, *p* = .02. In contrast, the individual empathy ratings did not correlate with the personal distress (PD) subscale of the IRI and the individual empathy ratings, *r*_(29)_ = -.04, *p* = .82. According to these results, the induced motive is related to empathic concern rather than personal distress.

### The financial incentive increased the frequency of prosocial decisions, in particular if empathy was low

Comparing the reaction times of prosocial decisions in the empathy-bonus and the empathy-alone condition revealed no significant difference, (lmm *χ^2^_(1)_ = 2.24*, *p* = .13, *B* = 0.27, *s.e.* = 0.18). There was also no difference when only selfish decisions were considered (lmm *χ^2^_(1)_ = 0.14*, *p* = .71, *B* = -0.08, *s.e.* = 0.22) and when all decisions were included (lmm *χ^2^_(1)_ = 1.99*, *p* = .16, *B* = 0.26, *s.e.* = 0.19).

The frequency of prosocial decisions was significantly higher in the empathy-bonus condition compared to the empathy-alone condition (**Fig. 3A**), (lmm *χ^2^_(1)_ = 14.35*, *p* < .01, *B* = - 0.57, *s.e.* = 0.15, *R²_m_* = .08). We also computed the percent change in prosocial decisions in the empathy-bonus condition relative to the empathy-alone condition ((empathy-bonus - empathy-alone)/empathy-alone * 100). The results revealed a significant relative increase of 23.88% (*s.e.* = 7.91%), *t_(30)_* = 3.02, *p* < .01.

**Fig. 3.**
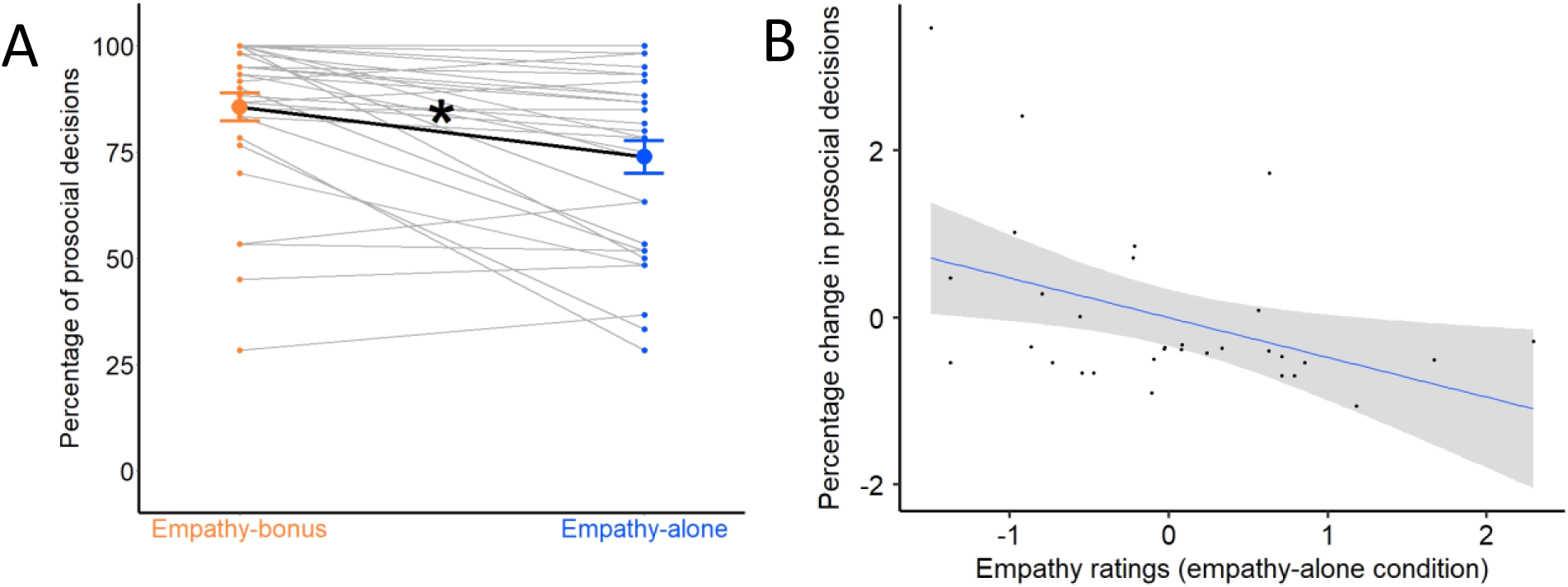
Percentage of prosocial decisions, reaction times and the relationship between the relative increase in prosocial decisions in the empathy-bonus condition and empathy ratings. **A)** Individual percentage of prosocial decisions in the empathy-bonus (orange) and the empathy-alone condition (blue). **B)** Negative relationship between the relative increase in prosocial decisions in the empathy-bonus condition and empathy ratings. The lower a participant’s empathy rating, the higher the incentive-related increase in prosocial decisions.

In an additional analysis, we compared the number of prosocial decisions in the empathy-alone condition with the number of prosocial decisions in a baseline condition (without any motive induction) from a previous study using a similar paradigm and the same allocation task (Hein *et al*., 2016b). The results revealed significantly more prosocial decisions in the empathy-alone condition compared to the baseline condition, empathy-alone (*M* = 73.92%, *s.e.* = 0.39), baseline condition (*M* = 49.37%, *s.e.* = 0.32), (*t_(59.963)_* = 4.85, p < .01).

We tested if that was induced before the decision task. A linear mixed model with reaction times of the prosocial decisions as dependent variable and empathy ratings, condition (empathy-alone / empathy-bonus) and empathy ratings × condition as predictors revealed a significant negative effect of empathy ratings (lmm *χ^2^_(1)_* = 6.61, *p* = .01, *B* = -0.36, *s.e.* = 0.17), which was comparable in both conditions, condition (lmm *χ^2^_(1)_* = 2.17, *p* = .14, *B* = 0.27, *s.e.* = 0.18), condition x empathy rating interaction (lmm *χ^2^_(1)_* = 0.02, *p* = .89, *B* = -0.02, *s.e.* = 0.18; *R²_m_* = .15). According to these results, higher empathy ratings predicted faster prosocial decisions.

A regression analysis with the percentage change in prosocial decisions as dependent variable and empathy ratings as predictor revealed a significant negative relationship (*B* = -0.42, *s.e.* = 0.17, *p* = .02, *R²* = .18). The lower an individual’s empathy ratings, the stronger the increase in the frequency of prosocial decisions in the empathy-bonus condition relative to the empathy-alone condition (**Fig. 3B**).

### The financial incentive increased the speed of information accumulation, but not the initial decision preference

To specify which component of the prosocial decision process was enhanced by the financial incentive, relative to prosocial decisions in the empathy-alone condition, we used hierarchical drift-diffusion modelling (HDDM; (Vandekerckhove *et al*., 2011; Wiecki *et al*., 2013)), a version of the classical drift-diffusion model that exploits between-subject and within-subject variability using Bayesian parameter estimation methods. We estimated the three aforementioned DDM parameters (*v*, *z*, *a*) for every condition and participant. Comparing the observed data with 500 datasets simulated by the HDDM (Wiecki *et al*., 2013) showed that the HDDM fit the data with 95% credibility (see quantile comparison table at github.com (https://github.com/Vassil-Iotzov/empathy_incentives).

We compared the speed of information accumulation (drift rate; *v*-parameters), the initial prosocial decision preferences (starting point; *z*-parameters), and the amount of integrated information (*a*-parameters) between the empathy-bonus and the empathy-alone condition. The comparison of the posteriors (Wiecki *et al*., 2013) revealed high probability for a larger *v*-parameter in the empathy-bonus condition compared to the empathy-alone condition, *v*-empathy-bonus (*M* = 2.03, *s.e.* = 0.22), *v*-empathy-alone (*M* = 1.24, *s.e.* = 0.19), (*p_(v-empathy-bonus > v-empathy-alone)_* = .99; **Fig. 4A**). In contrast, the probability for a differences between the other decision parameters was relatively low, *z*-empathy-bonus (*M* = 0.47, *s.e.* = 0.01), *z*-empathy-alone (*M* = 0.46, *s.e.* = 0.01; *p_(z-empathy-bonus > z-empathy-alone)_* = .54), *a*-empathy-bonus (*M* = 1.96, *s.e.* = 0.08), *a*-empathy-alone (*M* = 1.88, *s.e.* = 0.09; *p_(a-empathy-bonus > a-empathy-alone)_* = .79). This indicates that financial incentives enhanced the efficiency of the prosocial decision process, while leaving initial prosocial preferences unchanged.

**Fig. 4.**
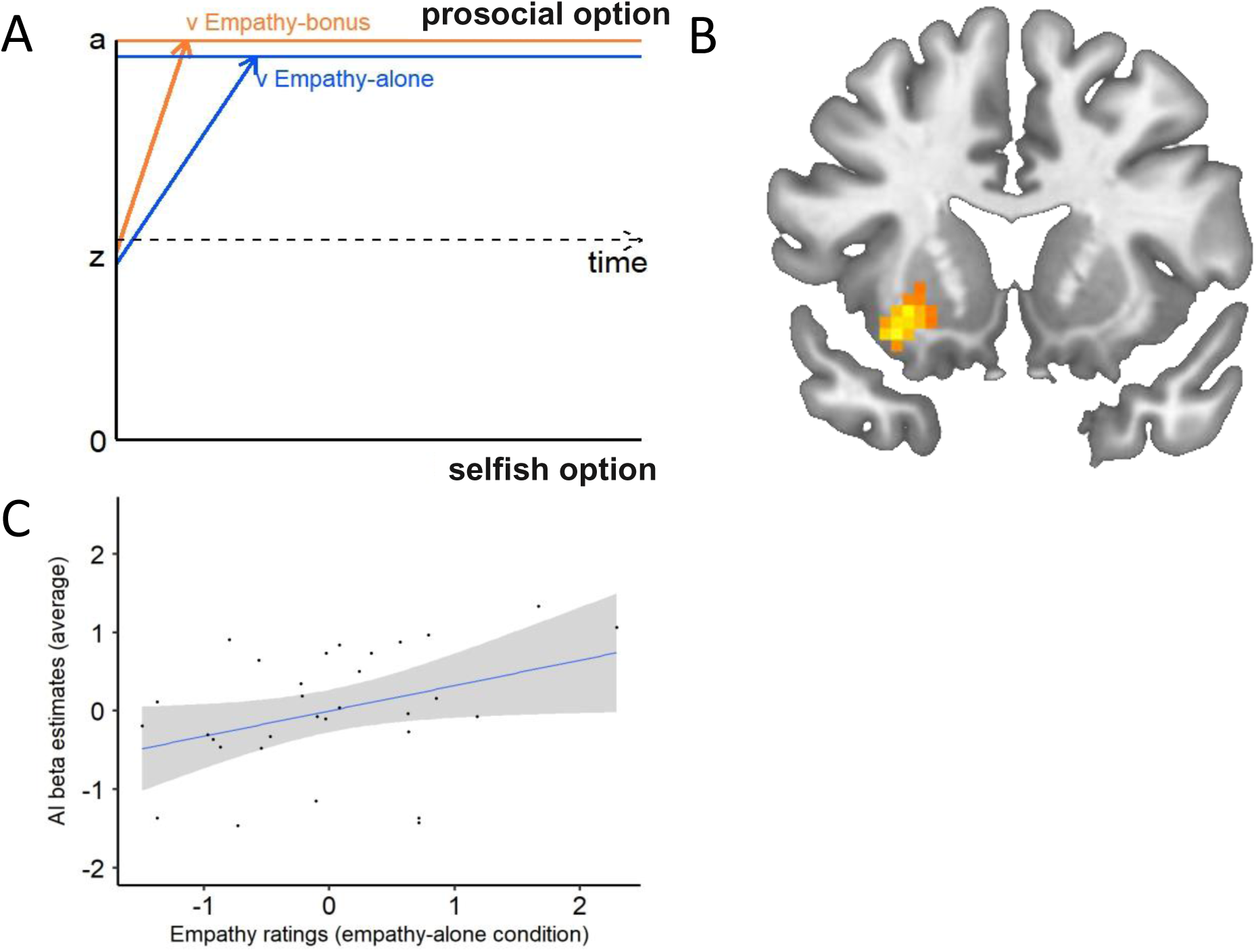
Drift-diffusion modelling (DDM) results and their relationship with neural responses in anterior insular cortex and empathy ratings. **A)** Visualization of the obtained DDM parameters showing an enhanced speed of information accumulation (*v*-parameter) in the empathy-bonus condition (orange) compared to the empathy-alone condition (blue). **B)** The neural response in the anterior insula (AI) correlates with the individual *v*-parameters in the empathy-bonus condition (visualized using 5% FWE cluster-based inference with P < .001 cluster-forming threshold; k = 50). The higher the speed of information accumulation in the empathy-bonus condition, the stronger the neural response in AI. **C)** Significant positive relationship between the individual strength of the AI response and the individual empathy ratings. The beta estimates reflect the average of AI activation from the empathy-bonus and the empathy-alone condition, extracted from the same AI clusters that correlated with the *v*-parameter in the empathy-bonus condition (shown in B).

Inspired by previous studies (Hutcherson *et al*., 2015; Chen and Krajbich, 2018), in an additional analysis, we conducted a model that took the trial-by-trials difference in points for self vs other into account. To do so, we added the point difference (point for self vs points for other) as additional covariate effecting the drift rate (Chen and Krajbich, 2018). The results replicated the observed findings (high probability for a larger *v*-parameter in the empathy-bonus condition compared to the empathy-alone condition: *v*-empathy-bonus (*M* = 5.69, *s.e.* = 0.22), *v*-empathy-alone (*M* = 4.94, *s.e.* = 0.19), *p_(v-empathy-bonus > v-empathy-alone)_* = .99), no differences between the other decision parameters *z*-parameter: *z*-empathy-bonus (*M* = 0.49, *s.e.* = 0.01), *z*-empathy-alone (*M* = 0.47, *s.e.* = 0.01), *p_(z-empathy-bonus > z-empathy-alone)_* = .70; *a*-parameter: *a*-empathy-bonus (*M* = 1.97, *s.e.* = 0.08), *a*-empathy-alone (*M* = 1.89, *s.e.* = 0.08), *p_(a-empathy-bonus > a-empathy-alone)_* = .69).

### The incentive-related facilitation of prosocial decisions and individual differences in empathy are associated with changes in anterior insula activation

First, we conducted the main contrasts between the prosocial decision-related activation in the empathy-bonus vs the empathy-alone conditions and vice versa. Based on the applied statistical threshold (P_(FWEvoxel-based)_ < .05) there were no significant results. This indicates that on average the same neural circuitries are involved in computing prosocial decisions driven by empathy and by empathy and the bonus.

Second, we identified neural regions that are related to an increase in drift rate in the empathy-bonus condition, i.e., the choice parameter that accounted for the facilitation of the prosocial decision process in the empathy-bonus compared to the empathy-alone condition. We regressed the individual *v*-parameters against the neural activation during prosocial decisions in the empathy-bonus condition, using a second-level regression. The results showed significant activation in the left anterior insula (MNI peak coordinates, x = -27, y = 38, z = 5; **Fig. 4B**) and the right lingual gyrus (MNI peak coordinates, x = 24, y = -67, z = -1, P(FWEvoxel-based < .05).

Exploratory analysis (5% FWE cluster-based inference with a cluster-forming threshold of P_uncorrected_ < .001) further revealed activations in the right AI, inferior lingual gyrus and pallidum (**Table 1**).

**Table 1.** Neural results of the second-level regression between prosocial decision-related activity in the Empathy-bonus condition and the speed of information accumulation (v-parameter) in the Empathy-bonus condition. The asterisk (*) indicates activations that are significant at 5% FWE voxel-based inference. We also conducted explorative analyses with 5% FWE cluster-based inference with a cluster-forming threshold of P < .001and a minimal cluster size of k = 50. Please note that peak-coordinates derived from cluster-wise inference only provide information about activated brain components, but not the exact brain region (Woo et al., 2014; Eklund et al., 2016).

| Region | Hemisphere | x y z | Cluster size | $t$-value | P(FWE<sub>cluster-based</sub>) |
| --- | --- | --- | --- | --- | --- |
| Anterior Insula | Left | -27 38 5 | 97 | 6.24 | .007* |
|  | Left | -30 14 -13 | 59 | 4.86 | .048 |
| Lingual gyrus | Right | 24 -67 -1 | 384 | 5.95 | .000* |
| Inferior lingual gyrus | Left | -51 -58 -19 | 181 | 5.23 | .000 |
| Pallidum | Left | -18 -7 -1 | 66 | 4.64 | .032 |

Third, inspired by previous evidence relating individual differences in AI responses to individual differences in empathy (Hein *et al*., 2010; Lamm *et al*., 2011; Marsh, 2018), we tested if the observed AI region (i.e., the region that correlated with the speed of information accumulation in the empathy-bonus condition) was also related to the empathy ratings that we collected prior to the allocation task. To do so, we extracted the average of the beta estimates related to prosocial decisions in the empathy-bonus and the empathy-alone condition from the entire activated AI clusters and regressed them against the individual differences in empathy ratings. The results showed a significant positive effect of empathy ratings (*B* = 0.41, *s.e.* = 0.19, *p* =.04, *R²* = .14). Because we used the average of the beta estimates from AI across both conditions, we can infer that the observed AI activation, in general processes individual differences in empathy, i.e., unbiased by the specific experimental conditions. The higher a participant’s empathy ratings, the stronger the neural response in the AI region, i.e. the same region that correlated with the speed of information accumulation in the empathy-bonus condition (**Fig. 4C**).

### The financial incentive has a differential effect on anterior insular activation in high and low empathic individuals

Given that the *v*-parameter and empathy ratings both are processed in the same AI region, it is plausible to assume that the two variables interact. To test that we conducted a linear mixed model with the beta estimates of AI activation during prosocial decisions in the empathy-bonus and the empathy-alone condition as a dependent variable. The individual *v*-parameters and empathy ratings were added as predictors, condition (empathy-bonus / empathy-alone) was added as a categorical variable. The results revealed significant main effects of condition (lmm *χ^2^_(1)_* = 12.26, *p* < .01, *B* = 0.67, *s.e.* = 0.19), empathy ratings (lmm *χ^2^_(1)_* = 4.43, *p* = .04, *B* = 0.33, *s.e.* = 0.16) and the *v*-parameter (lmm *χ^2^_(1)_* = 25.60, *p* < .01, *B* = 0.68, *s.e.* = 0.13). Moreover, there were significant interactions between empathy ratings x *v*-parameter (lmm *χ^2^_(1)_* = 5.60, *p* = .02, *B* = -0.40, *s.e.* = 0.17), and condition x *v*-parameter (lmm *χ^2^_(1)_* = 4.23, *p* = .04, *B* = -0.41, *s.e.* = 0.20), but not between condition x empathy ratings (lmm *χ^2^_(1)_* = 0.14, *p* = .71, *B* = 0.08, *s.e.* = 0.22). Finally, the analysis showed a significant condition x *v*-parameter x empathy rating interaction (lmm *χ^2^_(1)_* = 10.75, *p* < .01, *B* = 0.70, *s.e.* = 0.21, *R²_m_* = .49).

**Table 1** shows that also other brain regions correlated with the individual increase in *v*-parameters in the empathy-bonus condition. To test if these regions are also shaped by the interaction between the empathy ratings and the *v*-parameter, we conducted the same analysis with the beta estimates extracted from the pallidum, right lingual gyrus and left inferior lingual gyrus. The results revealed no significant interactions between empathy ratings and the *v*-parameter and no significant empathy ratings x *v*-parameter x condition interactions in any of these regions (empathy ratings x *v*-parameter, pallidum (lmm *χ^2^_(1)_* = 1.53, *p* = .22, *B* = -0.27, *s.e.* = 0.21), right lingual gyrus (lmm *χ^2^_(1)_* = 0.91, *p* = .34, *B* = -0.20, *s.e.* = 0.21), left inferior lingual gyrus (lmm *χ^2^_(1)_* < 0.01, *p* = .98, *B* = -0.01, *s.e.* = 0.22); empathy ratings x *v*-parameter x condition, pallidum (lmm *χ^2^_(1)_* = 0.77, *p* = .38, *B* = 0.24, *s.e.* = 0.27), right lingual gyrus (lmm *χ^2^_(1)_* = 0.48, *p* = .49, *B* = 0.19, *s.e.* = 0.28), left inferior lingual gyrus (lmm *χ^2^_(1)_* = 0.22, *p* = .64, *B* = 0.13, *s.e.* = 0.29)). This indicates that the observed effects are specifically related to neural responses in the AI.

To unpack the significant condition x *v*-parameter x empathy rating interaction in AI, we tested the relationship between the *v*-parameter and the empathy ratings separately in the empathy-alone and the empathy-bonus condition. We found a significant negative empathy x *v*-parameter interaction in the empathy-bonus condition (*B* = -0.37, *s.e.* = 0.14, *p* = .02), with significant main effects of *v* (*B* = 0.70, *s.e.* = 0.11, *p* < .01) and empathy ratings (*B* = 0.29, *s.e.* = 0.13, *p* = .04, *R²* = .65; **Fig. 5A**). The results for the empathy-alone condition revealed a marginal significant positive empathy x *v*-parameter interaction (*B* = 0.30, *s.e.* = 0.16, *p* = .07) with a significant main effect of the empathy ratings (*B* = 0.43, *s.e.* = 0.18 *p* = .03) and no main effect of the *v*-parameter (*B* = 0.25, *s.e.* = 0.18, *p* = .19; *R²* = .31, **Fig. 5B**).

**Fig. 5.**
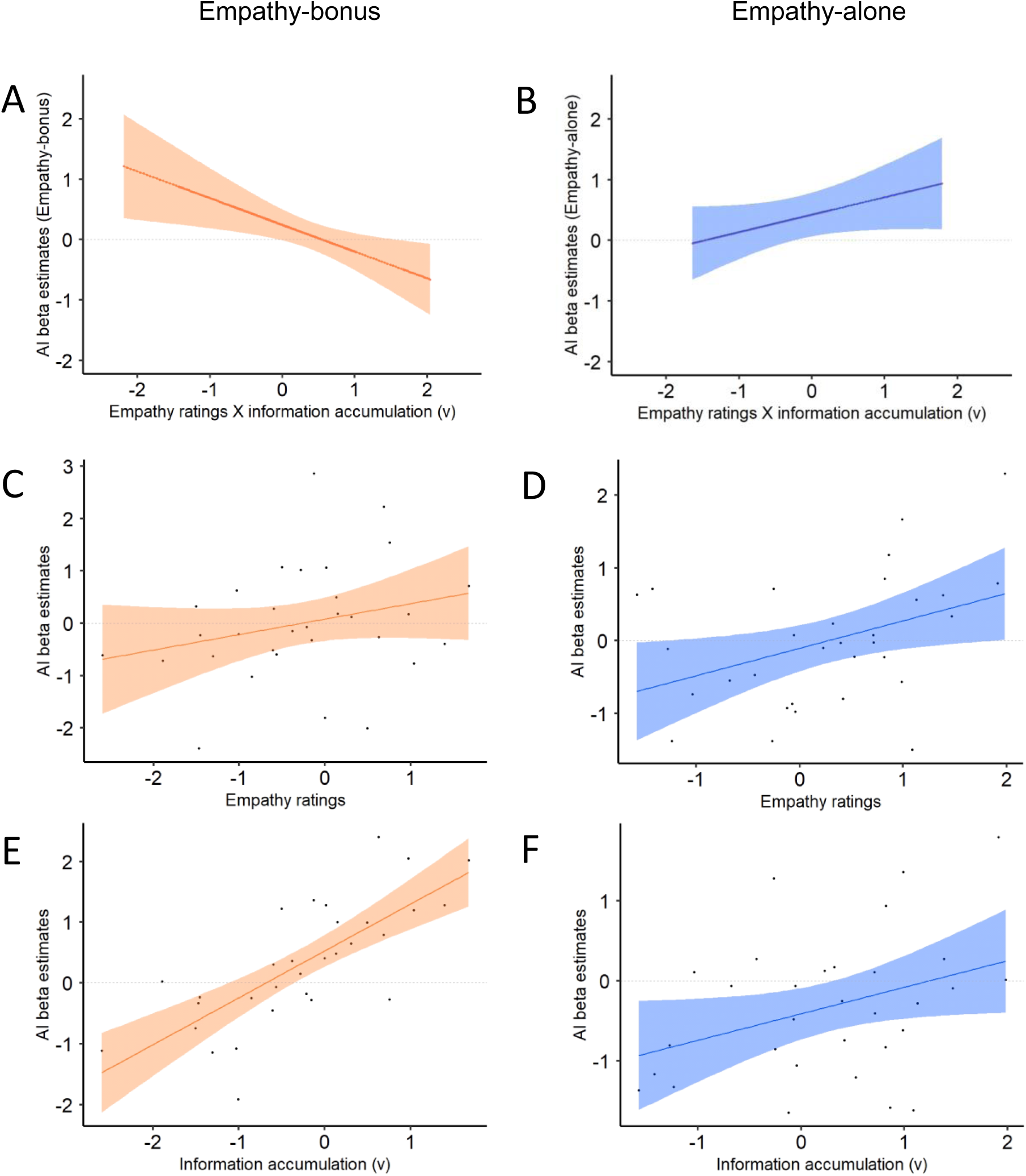
Relationships between anterior insula (AI) beta estimates and empathy ratings and between AI beta estimates and speed of information processing (*v*-parameter) in the empathy-bonus and empathy-alone conditions. The beta estimates reflect the average of AI activation from the empathy-bonus and the empathy-alone condition, extracted from the same AI clusters that correlated with the *v*-parameter in the empathy-bonus condition (shown in Fig. 4B). **A)** Effect of the empathy ratings x *v*-parameters interaction on AI responses in the empathy-bonus condition. **B)** Effect of the empathy ratings x *v*-parameters interaction on AI responses in the empathy-alone condition. **C)** The relationship between the individual strength of the AI responses and the individual empathy ratings in the empathy-bonus condition was not significant. **D)** Significant positive relationship between the individual strength of the AI responses and the individual empathy ratings in the empathy-alone condition. **E)** Significant positive relationship between the individual strength of the AI responses and the speed of information processing (*v*-parameter) in the empathy-bonus condition. **F)** Significant positive relationship between the individual strength of the AI responses and the speed of information processing (*v*-parameter) in the empathy-alone condition.

To further unpack the two-way interactions, we tested the relationship between the *v*-parameter and anterior insula (AI) beta estimates, as well as the relationship between empathy ratings and AI beta estimates separately in the empathy-bonus and the empathy-alone condition. Given that empathy facilitates prosocial decisions (Batson *et al*., 1995; Decety *et al*., 2016) and correlates with neural responses in AI cortex, we assumed a positive relationship between the empathy ratings and the drift ratings and empathy ratings and AI activation. To test these apriori assumptions, we used one-sided tests (Pfaffenberger and Patterson, 1977; Ruxton and Neuhäuser, 2010). In the empathy-alone condition, the results revealed a significant positive relationship between *v*-parameter and AI beta estimates (*B* = 0.38, *s.e.* = 0.19, *p* = .02, **Fig. 5F**), a significant positive relationship between empathy ratings and AI beta estimates (*B* = 0.43, *s.e.* = 0.18, *p* = .01, **Fig. 5D**), and a significant positive relationship between empathy ratings and drift rate (*B* = 0.30, *s.e.* = 0.18, *p* = .05). In the empathy-bonus condition we observed a significant positive relationship between *v*-parameter and AI beta estimates (*B* = 0.73, *s.e.* = 0.12, *p* < .01, **Fig. 5E**), while the relationships between empathy ratings and AI beta estimates (*B* = 0.23, *s.e.* = 0.16, *p* = .08, **Fig. 5C**) and between empathy ratings and drift rate were not significant (*B* = 0.21, *s.e.* = 0.17, *p* = .10). The finding of a positive relationship between empathy ratings and the drift rate and empathy ratings and AI beta estimates in the empathy-alone condition is in line with previous evidence showing that empathy facilitates prosocial decisions (Batson *et al*., 1995; Decety *et al*., 2016). In the empathy-bonus condition, the relationship between empathy ratings and drift rate and empathy ratings and AI estimates was no longer significant, indicating that in the presence of an incentive, empathy was no longer a significant driver of prosocial decisions. Interestingly, the interaction between the empathy ratings and the drift rate reduced AI activation in the empathy-bonus condition while increasing it in the empathy-alone condition. This indicates that in the empathy-bonus condition the empathy ratings (indicating the strength of the empathy motive before the bonus was offered) suppress the positive effect of the *v*-parameter on the neural response in AI.

To test the robustness of the differential effects in the empathy-bonus and the empathy-alone conditions, we extracted the beta-estimates of prosocial decision-related activation in the empathy-bonus and the empathy-alone condition from an independent region of interest in the AI (defined based on the peak coordinates reported in a recent meta-analysis on empathy of pain studies (Jauniaux *et al*., 2019). We conducted a linear mixed model with these beta-estimates as dependent variable, and condition (empathy-bonus / empathy – alone), empathy ratings, and *v*-parameters as predictors. The results replicated the significant condition x *v*-parameter x empathy rating interaction reported above (lmm *χ^2^_(1)_* = 5.81, *p* = .02, *B* = 0.61, *s.e.* = 0.25, *R²_m_* = .19), reflecting a negative relationship in the empathy-bonus condition and a positive relationship in the empathy-alone condition.

## Discussion

Our study investigated how financial incentives affect empathy-related prosocial decisions. The results show that on average financial incentives increase the frequency of prosocial decisions (**Fig. 3A**), in particular in individuals that scored low on empathy (**Fig. 3B**). The finding that the financial bonus enhanced the frequency of prosocial decisions is in line with previous studies showing an incentive-related increase in prosocial behaviours (Balliet *et al*., 2011; Stoop *et al*., 2018). Extending this previous evidence, our results reveal that this effect is modulated by individual differences in empathy, i.e., stronger if a person’s empathic motivation is low. Besides providing insights into the interplay between financial incentives and empathy, our results specified how financial incentives affect the prosocial decision process. The results of drift-diffusion modelling showed that the financial incentive enhanced the efficiency (i.e., speed of information accumulation captured by the *v*-parameter) of prosocial decisions in the empathy-bonus compared to the empathy-alone condition (**Fig. 4A**). In contrast, the incentive had no significant effect on participants’ initial prosocial preferences, i.e., the preference of making a selfish or prosocial decision with which they entered the decision process (captured by the *z*-parameter).

Outside the domain of prosocial decisions, there is evidence that the efficiency of decisions (captured by the *v*-parameter) is affected by individual differences in emotions (Lerche *et al*., 2018; Roberts and Hutcherson, 2019; Aylward *et al*., 2020; Thompson and Steinbeis, 2021). For example, according to the results of Thompson and Steinbeis (2021), individuals with greater state anxiety show increased *v*-parameter on fearful face trials. Extending these findings, our results reveal that the speed of information accumulation is shaped by the motivation that drive participants’ prosocial decisions, i.e., higher if a prosocial decision is rewarded than if it is only based on empathy.

On the neural level, the incentive-related facilitation of the prosocial decision process was related to the participants’ neural response in the left anterior insula (AI; **Fig. 4B**). Previous neuroscience research has associated the anterior insula activity with empathy (Hein *et al*., 2010; Lamm *et al*., 2011; Masten *et al*., 2011; Hein *et al*., 2016b; Marsh, 2018) and the propensity for prosocial decisions (Hein *et al*., 2010; Masten *et al*., 2011; Hein *et al*., 2016b; Marsh, 2018). In line with this previous evidence, our results show that the facilitation of prosocial decisions (captured by an increased speed of information accumulation) is related to an increase of AI responses (**Fig. 4B**) and that this same AI region also correlated with individual differences in empathy (**Fig. 4C**).

Adding a novel aspect, our findings reveal how financial incentives alter the effect of empathy on the computation of prosocial decisions in the anterior insular cortex. After offering a bonus in the empathy-bonus condition, the relationship between empathy ratings and drift rate and empathy ratings and AI estimates was no longer significant, indicating that in the presence of an incentive, empathy was no longer a significant driver of prosocial decisions. Interestingly, the interaction between the empathy ratings and the drift rate significantly reduced AI activation in the empathy-bonus condition (**Fig. 5A**) while increasing it in the empathy-alone condition (**Fig. 5B**). This indicates that in the empathy-bonus condition, the strength of the empathy motive (captured by the individual strength of the empathy ratings before the bonus was offered) suppressed the positive relationship between information accumulation during prosocial decisions and the neural response in AI. Together, these findings indicate that the anterior insula integrates self-regarding (gaining the financial incentive) and other-regarding (empathy with the other person) motives that both elicit prosocial decisions and thus forms a plausible neural basis for the impact of financial incentives on empathic motivation.

In our study, empathy was conceptualized as a motive that can drive prosocial decisions. And indeed, the empathy ratings of our participants that correlated with empathic concern (but not personal distress) facilitated the prosocial decision process in the empathy-alone condition, in line with previous findings (Batson *et al*., 1995; Decety *et al*., 2016). That said, the result that financial incentives counteracted the facilitating effect of empathy on prosocial decisions in highly empathic individuals might indicate that highly empathic individuals are less motivated to empathize in the presence of an incentive, an assumption that supports the notion that empathy itself is a motivated state (Zaki, 2014).

The financial incentive for prosocial decisions was offered in private, and self-image concerns were reduced as far as possible, at least with regard to public reputation. However, some highly empathic participants nevertheless showed an incentive-related decline in prosocial decisions (see also **Fig. 3B**). It is conceivable that highly empathic participants feel insulted by the bonus because “being paid to be nice” undermined their intrinsic empathic motivation that otherwise (i.e., in the empathy-alone condition) drives their prosocial decisions. Thus, although on average our findings show that the incentive increased the frequency of prosocial decisions compared to an empathy-alone condition, it is still possible that it undermines prosocial behavior in highly empathic participants. To test this assumption, future studies should test the effect of financial incentives on empathy-based decisions in extreme groups, i.e., groups of extremely high or low empathic individuals. Moreover, it would be interesting to use a trial-by-trial bonus manipulation that allows for modelling the effect directly as part of the DDM.

In summary, our current results indicate that financial incentives offered in private facilitate prosocial decisions in low empathic individuals but have little effect in case of strong empathic motivation.

## Acknowledgments

We thank Isabelle Ehrlich for help with data collection and Marthe Gründahl and Martin Weiß for feedback on the manuscript.

## Authors’ Contributions

Grit Hein and Vassil Iotzov designed the research with input from Jochen Kaiser; Vassil Iotzov programmed the experiment with input from Anne Saulin and performed the research; Vassil Iotzov and Anne Saulin analyzed the data with input from Grit Hein, Jochen Kaiser and Shihui Han; Grit Hein and Vassil Iotzov wrote the paper with input from Anne Saulin, Jochen Kaiser and Shihui Han.

